# Physiologically relevant metrics for evaluating photoreceptor signal reproduction in visual displays

**DOI:** 10.1101/2021.02.27.433203

**Authors:** Allie C. Schneider, Takuma Morimoto, Hannah E. Smithson, Manuel Spitschan

## Abstract

The colour gamut of a display defines the range of colour signals that it can reproduce. For human vision this range is typically visualized in a colour space that reflects the spectral sensitivities of the cone photoreceptors. While the cones dominate “canonical” visual function (i.e. perception of colour, form, and motion) under photopic light levels, rods and melanopsin-containing intrinsically photosensitive retinal ganglion cells (ipRGCs) also respond to light. Three-primary display technologies, with their focus on reproducing colour, are not designed to control the rod and melanopsin excitations. As a result, conventional display metrics do not capture the display’s ability to reproduce the full human physiological response to light. In this paper, two physiologically relevant metrics are proposed for quantifying the reproduction and distortion of the photoreceptor signals by visual displays. An equal-luminance photoreceptor excitation diagram is proposed, extending the MacLeod-Boynton chromaticity diagram, providing three-dimensional projections of the five-dimensional photoreceptor signal space to help with visualization. Each metric and visualisation quantifies a specific aspect of the reproducibility of real-world spectra, analogous to the colour rendering index. To demonstrate their application, we use these tools to assess both real and hypothetical displays across tens of thousands of radiance spectra derived from real-world reflectance samples under various illuminants.

## 1. Introduction

As reliance on display technology grows, manufacturers aim to create hyper-realistic displays that evoke the full human physiological response to real-world light. This response can be represented in a five-dimensional sensor space defined by the excitations of the three cone types, rods, and melanopsin-containing intrinsically photosensitive retinal ganglion cells (ipRGCs). These spectral sensitivities are standardized in CIE S026 [1], and the term α-opic denotes the response magnitude of each photoreceptor. However, no existing metrics characterize a display’s ability to reproduce the pattern of α-opic activations—referred to as the “α-opic quintuplet.” As display technology evolves to replicate rod and melanopsin excitations, suitable metrics and visualization tools will be essential. This paper introduces physiologically relevant metrics to evaluate α-opic reproduction and proposes an equal-luminance photoreceptor excitation diagram, extending the MacLeod–Boynton chromaticity diagram [2], to visualize the full photoreceptor space.

### 1.1. The human physiological response to light

The need to quantify the α-opic quintuplet for display characterization arises from the physiology of the human light response. This response begins with photon absorption by retinal photoreceptors. In photopic vision, light is detected by three cone types: long- (L), medium- (M), and short-wavelength-sensitive (S) cones. Post-receptoral neurons with center-surround receptive fields support spatial pre-processing, while comparisons between cones with different spectral sensitivities enable chromatic pre-processing.

The cones share the role of light sensation with rods and ipRGCs, making the visual response to light more complex than the cone-driven photopic model alone. Rods are well known for their role in low-light vision [3,4], but their saturation point in bright conditions remains poorly defined and varies across individuals and viewing contexts. Recent studies suggest that rod activity persists well into the photopic range [5–9]. Even under the classical saturation model, typical office lighting falls within the mesopic range, indicating that rods likely contribute to everyday visual experience [10, 11]. Additionally, evidence suggests that ipRGC signals reach the visual cortex, with growing support for melanopsin’s role in visual functions such as brightness perception, contrast sensitivity, and color processing [9, 12–22].

Beyond vision, light plays a key role in entraining the internal circadian clock to the external light–dark cycle. Exposure to bright light in the evening or at night can suppress the production of melatonin, colloquially referred to as the “sleep hormone” [23–25]. Importantly, the spectral composition of light, not just its overall intensity, influences these circadian effects, with the spectral sensitivity of non-visual light responses closely matching that of melanopsin in the ipRGCs [26, 27]. Thus melanopsin excitation is often attributed as being the primary driver of these non-visual circadian effects of light. ipRGCs also receive synaptic input from rods and cones [28], and therefore these photoreceptors may also be contributing to the circadian effects of light.

ipRGCs work alongside the cones to drive other non-visual responses to light. For instance, pupil size is regulated by melanopsin and cone signals, and possibly by rods under certain conditions [14, 29–35]. L- and M-cone signals act synergistically with melanopsin to influence pupil size, while S-cone signals interact with melanopsin in an opposite manner [36, 37]. ipRGCs also contribute to light-induced alertness [38–40], and light exposure has been shown to affect mood, with light therapy used as a treatment for Seasonal Affective Disorder (SAD) and other mental health conditions [25, 41].

In sum, rods and ipRGCs play key roles in the physiological and behavioral effects of light. Beyond the cone photoreceptor, melanopsin may also contribute to visual function, while rods remain active even under conditions traditionally attributed solely to cones [6, 8, 9, 12–20]. These insights highlight the need to quantify rod and melanopsin signal reproduction in displays, beyond the scope of traditional colorimetry.

### 1.2. The reproduction of light by displays

Modern displays with wide color gamuts, high resolution, and high dynamic range can accurately reproduce the spatial and chromatic characteristics of real-world scenes. Consider reproducing a single chromatic sample, such as the radiance spectrum in Fig. 1(b), on a display with three primaries such as those in Fig. 1(c), rendered as a uniform color patch filling the entire field of view.

**Figure 1.**
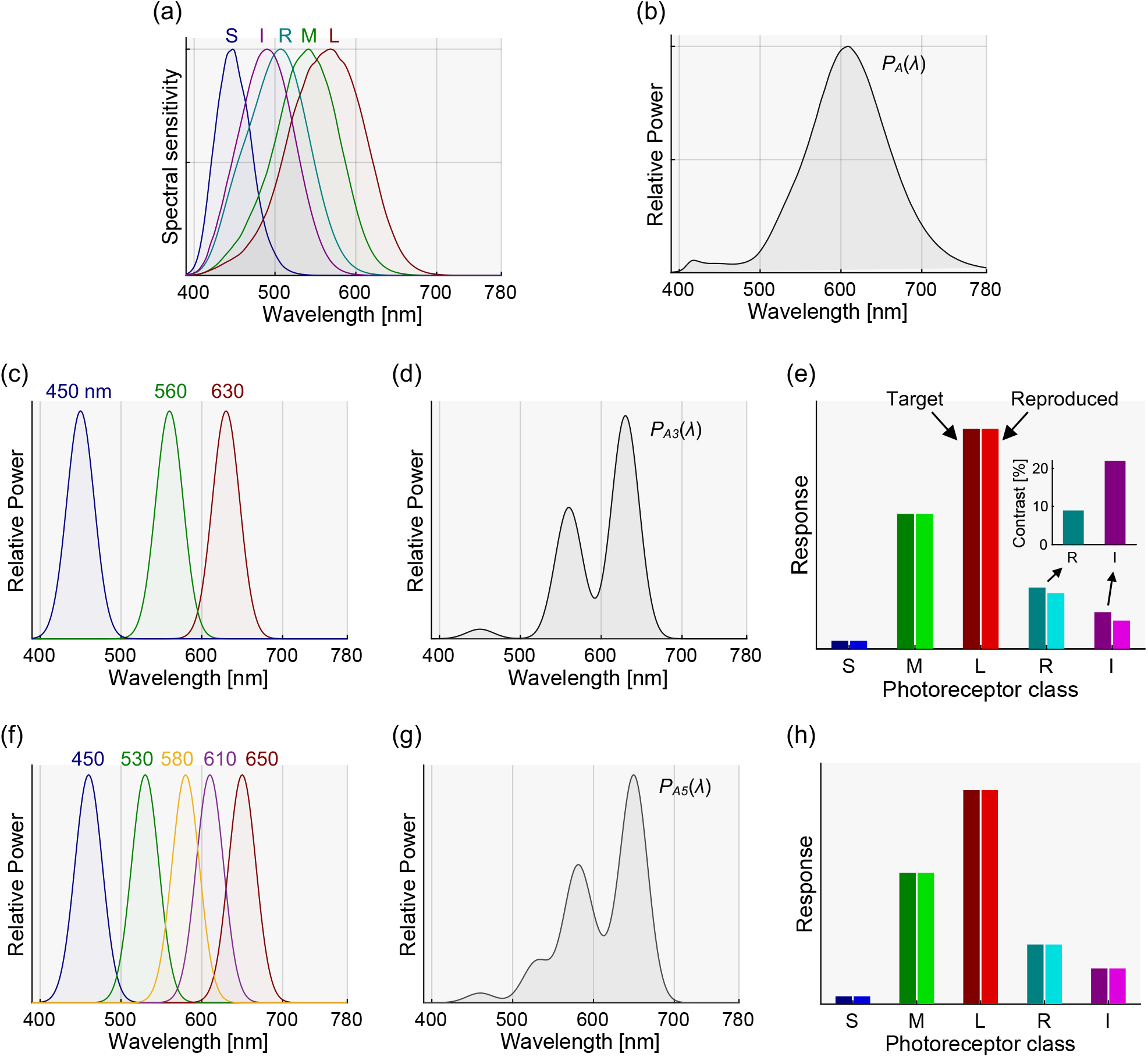
Examples illustrating how a display may reproduce and distort photoreceptor signals. (a) The spectral sensitivity functions, *T*_*k*_ (*λ*), of the cones (*k* ∈ [*L, M, S*]), rods (*k* = *R*), and melanopsin (*k* = *I*), as defined in the CIE S026 standard [1, 26, 42, 43]. (b) A radiance spectrum from a real-world surface reflectance viewed under a real-world illuminant *P*_*A*_(*λ*). (c) Example primaries, *p*(*λ*), of a hypothetical three-primary display. (d) The spectral output, *P*_*A3*_(*λ*), of the hypothetical three-primary display (c) that reproduces the same cone excitations as the real-world radiance spectrum in (b). (e) The cone, rod, and melanopsin excitations from the real-world radiance spectrum, *A* (the left hand bar in each pair, in darker colours) and from the hypothetical three-primary display’s spectral output, *A*_3_’ (the right hand bar in each pair in lighter colours). The inset shows the absolute percent contrast between the signals from the real-world spectrum and the display’s spectral output. (f) Example primaries, *p*(*λ*), of a hypothetical five-primary display. (g) The spectral output, *P*_A5_(*λ*), of the hypothetical five-primary display (f) that reproduces the same photoreceptor excitations as the real-world radiance spectrum (b). (h) The photoreceptor excitations from the real-world spectrum, *A*, (left bar in each pair, darker colours) and the hypothetical five-primary display’s spectral output, *A*_5_’, (right bar in each pair, lighter colours). While there is a mismatch in rod and melanopsin signals for the three-primary displays, the five-primary displays can accurately reproduce the α-opic quintuplet.

To match the display color to that of the real-world spectrum, one can compute a relative intensity vector, *C* = [*r, g, b*], such that 0 <= *r* <= 1, 0 <= *g* <= 1, and 0 <= *b* <= 1,with spectral power distribution for the display primaries *p*_*r*_(*λ*), *p*_*g*_(*λ*), and *p*_*b*_(*λ*). This vector *C* is chosen such that the resulting spectral output (Fig. 1(d)) elicits the target L, M, and S cone responses, *A* = [*A*_*L*_, *A*_*M*_, *A*_*S*_], derived from the real-world spectrum, assuming *A* lies within the display’s gamut and intensity range. These relationships can be written as Eq. 1, where *B* is the conversion matrix from *C* to *A*, and *B*^*-1*^ is its inverse.

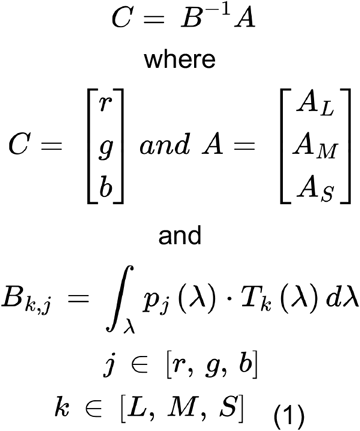

Note that in this example, the Stockman–Sharpe 10° cone fundamentals—*T*_*L*_(*λ*), *T*_*M*_(*λ*), *T*_S_(*λ*), —are used to calculate the L, M, and S responses. These fundamentals are defined in the CIE S026 standard and depicted in Fig. 1(a) [1, 42–44]. The L, M, and S responses from both the display *A*’ and the real-world spectrum *A* are equal, as shown in Fig. 1(e). However, the rod(*R*) and melanopsin(*I*) signals *A*_R_’ and *A*_I_’ remain uncontrolled and depend on the display’s spectral output *P*(*λ*), which is defined by the spectra of the primaries and the intensity vector *C* used to match the cone responses (Eq. 2). The distorted rod and melanopsin signals are computed by Eq. 3. The CIE S026 standard spectral sensitivity functions for rods *T*_*R*_ (*λ*), and melanopsin, *T*_*I*_ (*λ*), are used here (Fig. 1(a)) [1, 26, 45]. Figure 1(e) shows an example of how these signals are distorted when the example spectrum is rendered on the hypothetical three-primary display shown in Fig. 1(c).

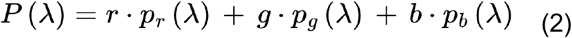

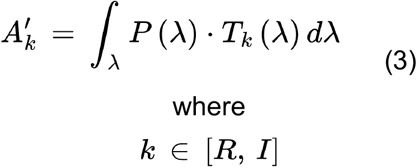

The inability of traditional displays to control rod and melanopsin excitation is becoming increasingly relevant, as screen use grows and technologies such as augmented and virtual reality lead to longer exposure in artificial, mesopic visual environments [46, 47]. Whether these signals are controlled or not may have a range of implications. Evening light exposure from devices may influence melanopsin-driven processes, potentially affecting melatonin levels, sleep, circadian rhythms, mood, and alertness [48–54]. Many displays offer a “night-shift” mode that reduces short-wavelength light by shifting the white balance toward longer wavelengths. However, this also alters the display’s chromaticity and visual appearance. Displays are often viewed under mesopic conditions, where rod activity can influence colour perception in both standard and “night-shift” modes [55–58]. Current “night-shift” approaches aim to reduce melanopsin activation by lowering short-wavelength light, though this may not fully address melatonin suppression [59]. While dimming may appear more effective than colour shifting, it also increases rod contribution due to reduced light levels. Beyond minimizing unintended melanopsin and rod activation, active control of these signals may be beneficial. For example, through tailored light exposure to support circadian alignment in shift workers [60]. Finally, the involvement of ipRGCs and rods in pupil size regulation may have consequences for spatial or 3D image reproduction. Challenges in replicating real-world focus cues have contributed to visual discomfort in some 3D display systems [61]. As pupil size is linked to accommodation, vergence, and depth of field, whether melanopsin-driven changes might influence these visual cues presents an interesting question for future investigation.

Throughout this work each evaluated spectrum is treated as a uniform, full-field stimulus: a single real-world radiance reproduced as a uniform patch filling the field of view, with the primary drive values solved independently for each spectrum to match the targeted photoreceptor signals. We do not model spatial structure, per-pixel constraints, or a shared backlight serving differently-driven pixels.

### 1.3. Towards more accurate physiologically relevant reproduction of light on displays

Advances in both hardware and software may improve a display’s ability to more accurately reproduce the α-opic quintuplet. Five-primary displays offer a way to control rod and melanopsin signals directly, without altering cone responses [31, 62–65]. With five primaries (denoted, *r, g, b, c, m*), it becomes possible to independently control all five photoreceptor signals [66], provided the desired signals fall within the display’s gamut, as described in Eq. 4 and illustrated in Fig. 1(f)–(h).

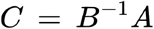

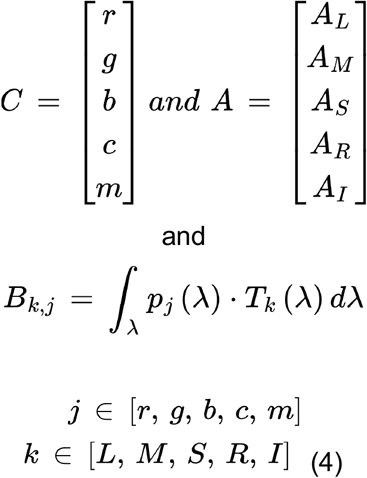

For most commercial displays that lack five primaries, is there an optimal approach to reproducing the α-opic quintuplet? In the case of three- and four-primary displays, it may be possible to design a more optimal chromaticity transformation that allows some controlled distortion in cone responses to reduce distortion in rod and melanopsin signals.

### 1.4. Quantifying a display’s physiologically relevant reproduction of light

To assess the control of rod and melanopsin responses in image reproduction systems, suitable metrics are needed to quantify their distortion and reproduction. Colour gamuts based on the CIE 1931 XYZ functions capture only chromaticity and photopic luminance (i.e., cone signals) [67]. The CIE 2015 (170-2) report adopts a more physiologically relevant colour space using the CIE 2006 (170-1) cone fundamentals, but it still quantifies only cone responses, excluding rods and melanopsin [44, 68].

We propose two physiologically relevant metrics to quantify the reproduction and distortion of photoreceptor signals on displays. The first, the Photoreceptor Signal Reproduction Metric (*PSRM*), measures how accurately a display can reproduce a set of photoreceptor signal quintuplets. The second, the Photoreceptor Signal Distortion Metric (*PSDM*), quantifies deviations between the displayed and intended photoreceptor signals. Additionally, an equal-luminance photoreceptor excitation diagram, an extension of the MacLeod-Boynton chromaticity diagram [2], is introduced to help visualize both real-world and reproduced photoreceptor signals.

## 2. Methods: two metrics and an equal-luminance photoreceptor excitation diagram

When designing metrics for α-opic reproduction, it is helpful to consider what constitutes ‘good’ performance. Two types of limitations can affect signal reproduction: (i) with fewer primaries than photoreceptor classes, independent control is not possible; and (ii) even with sufficient primaries, some target signals may lie outside the achievable gamut of the display. In chromaticity reproduction, all visible colours are bounded by the spectrum locus. However, three-primary displays, defined by a triangular gamut, cannot encompass the full range of achievable chromaticities. The sRGB, Adobe RGB, and Rec. 709 gamuts are widely used as standard references for evaluating chromaticity reproduction in three-primary displays [69–71]. In five-dimensional space, quantifying coverage of the available locus is more challenging. An alternative approach, used in colour rendering under various illuminants, is to define performance relative to a predefined reference set of spectra. This paper adopts that approach. The ideal reference set would include all real-world spectra, weighted by their likelihood of occurrence, but such a dataset does not yet exist [72]. However, existing datasets of real-world surfaces and illuminants can be combined to derive a set of radiance spectra that captures the diversity typically encountered.

The reference dataset used here comprises 18,216 radiance spectra, generated by combining 99 real-world reflectance samples with 184 reference illuminants. The reflectance samples are those developed for the IES TM-30 colour fidelity method [73] and represent natural materials, skin, textiles, paints, plastics, prints, and colour systems, with broad coverage of colour and wavelength space. The illuminants are selected from the CIE daylight series and the 401-illuminant set reported in [74], while the reflectance set is retained without modification. The final illuminant set comprises 65 CIE daylight phases spanning correlated colour temperatures from 4000 to 20,000 K in 250 K steps and 119 measured illuminants selected from the 401-illuminant set. The selected 119 illuminants satisfy the ANSI/IES TM-30-18 [75] thresholds of Rf ≥ 85 and Rg ≥ 90, consistent with the high-fidelity object-illumination guidance in Annex E. This screening retains realistic, high-fidelity sources and excludes spectra with poor colour-rendering performance. The resulting standardised dataset serves as a fixed benchmark for evaluating reproduction, rather than as a recommendation of particular illuminants. In this study, the term “real-world” spectra refer to measured natural and artificial illuminants and to reflectance spectra from physical surfaces.

The radiant spectra, *ψ*(*λ*), are computed from all possible combinations of reflectance spectra, *υ*(*λ*), and illuminant spectra, ι(*λ*), as described in Eq. 5, for a single case:

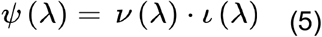

As the focus of this paper is on the reproduction of the α-opic quintuplet to within a scalar, the illuminant spectra were normalized to unit area, as in Eq. 6:

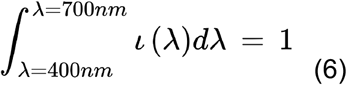

### 2.1. Metric I: Photoreceptor Signal Reproduction Metric (*PSRM*)

The first proposed metric, termed the Photoreceptor Signal Reproduction Metric (*PSRM*), quantifies the proportion of real-world α-opic quintuplets that a display can accurately reproduce. This metric is useful for comparing the extent of five-dimensional gamuts across different displays. It serves as a five-dimensional analogue to conventional gamut comparisons, such as those made with respect to the sRGB gamut. In traditional chromaticity assessments, the relative area within the chromaticity diagram that falls inside the display gamut is measured. Similarly, the *PSRM* evaluates the proportion of *n* real-world α-opic quintuplets, *A* = [*A*_L_, *A*_*M*_, *A*_S_, *A*_*R*_, *A*_*I*_], that can be reproduced on the display, *A*’ = [*A*_L_’, *A*_M_’,

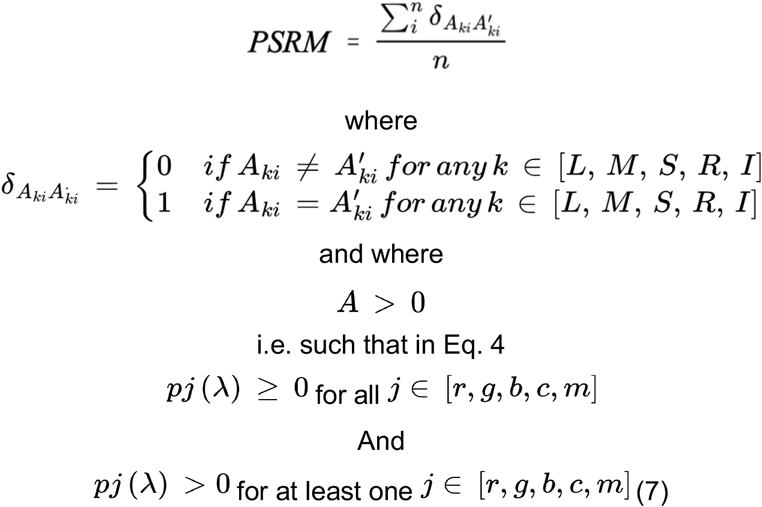

Note that it is not strictly required for the primary weights to be below a maximum of one, as any values exceeding this can be uniformly scaled to achieve reproduction. The denominator is the total number of spectra. The reproduced quintuplet [*A*_L_’, *A*_M_’, *A*_S_’, *A*_R_’, *A*_I_’] is counted only if it matches the corresponding real-world target [*A*_L_, *A*_*M*_, *A*_S_, *A*_*R*_, *A*_*I*_]. In other words, successful reproduction requires all five photoreceptor signals to satisfy the tolerance criterion and every primary weight to be non-negative, with at least one weight strictly positive. The *PSRM* ranges from 0 (no accurate reproduction) to 1 (perfect reproduction), or 0% to 100%.

In practice, some tolerance is required. Here, an α-opic signal for a given spectrum *i* is considered accurately reproduced (i.e. *A*_ki_’ = *A*_ki_) if the reproduced value for each photoreceptor is within 1% of error of the real-world signal as defined in Eq. 8.

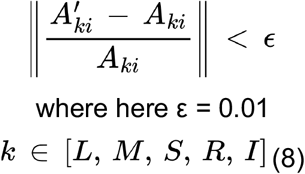

### 2.2. Metric II: Photoreceptor Signal Distortion Metric (*PSDM*)

The second metric, the Photoreceptor Signal Distortion Metric (*PSDM*), quantifies the distortion introduced into the uncontrolled α-opic signals. This metric captures the extent to which real-world photoreceptor signals are not accurately reproduced, helping to characterize both a display’s limitations and the potential impact of signal distortion on the observer. This metric is especially useful when displays have similar *PSRM* values, as it enables evaluation of distortions in specific photoreceptors of interest. For a given photoreceptor *k* and a particular spectrum *i*, the *PSDM*_*ki*_ is defined as the contrast between the reproduced signal *A*_*k*i_’, and the real-world target signal *A*_ki_, as shown in Eq. 9.

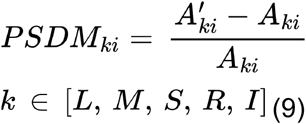

The *PSDM*_*ki*_ is calculated only for spectra that lie within the display’s reproducible gamut. Note that *PSDM*_*ki*_ corresponds to the same error metric used in the *PSRM* (Eq. 8).

While *PSDM*_*ki*_ can be computed for each spectrum individually, it is often useful to summarize distortion across photoreceptors and/or spectra. The average *PSDM*_*k*_ per photoreceptor *k* is defined as the mean of the absolute values of the *PSDM*_*ki*_ across all *n* spectra, as in Eq. 10. When signed medians are reported below, they refer to the distribution of the signed PSDMki values; PSDMk in Eq. 10 remains the mean absolute distortion.

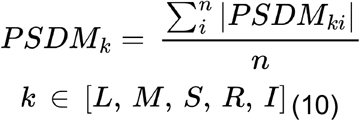

Furthermore, the overall *PSDM* for a display pooled across photoreceptors is defined as the mean absolute value of *PSDM*_*k*_, as shown in Eq. 11.

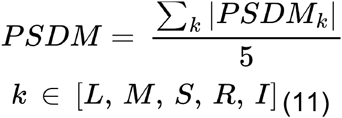

The minimum (and most desirable) value of *PSDM*_*k*_ for each photoreceptor *k*, and thus the overall *PSDM*, is zero, indicating no distortion in the reproduced signals. In theory, there is no fixed upper limit as the maximum *PSDM* depends on the specific display primaries and the considered real-world spectra. By design for current use-cases, three-primary display reproduce the cone signals (*PSDM*_*L*_ = *PSDM*_*M*_ = *PSDM*_*S*_ = 0). However, it is possible to define a transformation matrix (B in Eq. 4) that, for example, preserves the melanopsin signal instead of the M cone signal, resulting in zero melanopsin distortion while introducing distortion to the M cone. Alternatively, a transformation could be designed to balance distortion across all five photoreceptors.

Because the PSDM is defined per photoreceptor (PSDM_ki_ in Eq. 9 and PSDM_k_ in Eq. 10), the distortion of each receptor signal can be reported individually in addition to the overall PSDM (Eq. 11).

### 2.3. Equal-luminance photoreceptor excitation diagram

The CIE 1931 xy chromaticity diagram is widely used to visualize chromaticities. Although less common in the display industry, the MacLeod–Boynton chromaticity diagram [2] is frequently used in vision science to represent cone excitations at equal luminance. The traditional MacLeod-Boynton chromaticity diagram plots *s*_*MB*_(*λ*) against *l*_*MB*_(*λ*), where *s*_*MB*_(*λ*) and *l*_*MB*_(*λ*) are defined as in Eq. 12:

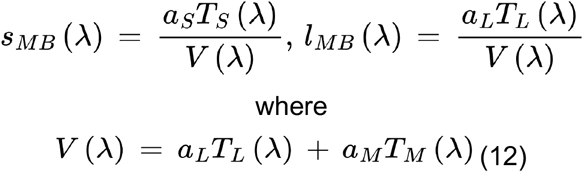

*V*(*λ*) and *s*_*MB*_(*λ*) are both normalized to peak at unity. The CIE 2006 10° spectral sensitivity functions, as defined in the CIE S026 standard, are also normalized to a unity peak [1, 42–44]. A wavelength range of 390 nm to 780 nm was used. When the spectral sensitivities are expressed in energy units, the scaling constants *a*_*S*_, *a*_*M*_, and *a*_*L*_ take the following values [2, 43]. The scaling constants used here are those specified for the CIE 2006 10° system in Eq. 7.6b of CIE 170-2; Eq. 7.4b applies to the corresponding 2° system.

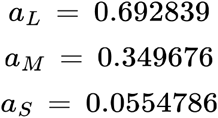

Fig. 2 (a) shows the spectrum locus in the traditional diagram. The MacLeod–Boynton space lends itself naturally to an extension that includes a melanopsin axis. Accordingly, a new equal-luminance photoreceptor excitation diagram is proposed, which plots the melanopic efficacy of radiance—that is, melanopsin excitation normalized by luminance [44, 68]. This new axis represents the variable *i*_*MB*_(*λ*), as defined in Eq. 13.

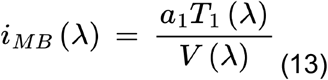

**Figure 2.**
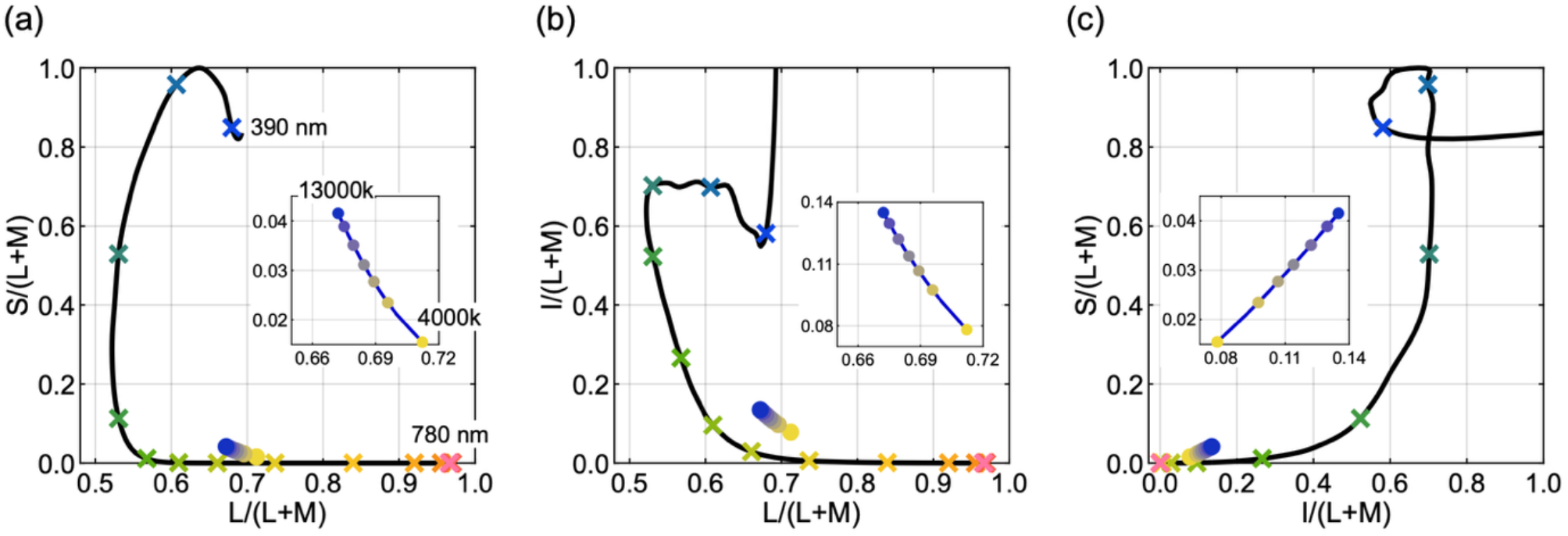
Proposed equal-luminance photoreceptor excitation space for representing melanopsin signals. (a) The spectrum locus in the MacLeod–Boynton chromaticity diagram (390–780 nm), with 25 nm steps marked by crosses and coloured to indicate wavelength progression. The daylight locus (4,000–13,000 K) is shown with a yellow-to-blue gradient. (b) Spectrum and daylight loci plotted as melanopic efficacy I / (L + M) vs. L / (L + M). (c) Alternative projection showing S / (L + M) vs. I / (L + M). Insets magnify the daylight locus in each plot.

The CIE S026 spectral sensitivity *T*_*I*_ (*λ*) function for melanopsin is used here, and normalised to unity peak [1, 26]. When the spectral sensitivities are expressed with respect to energy units, the scaling constant, *a*_*I*_, takes the following value in order for *i*_*MB*_(*λ*) to peak at unity:

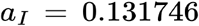

The spectral locus is shown in the proposed equal-luminance photoreceptor excitation diagram in Fig. 2(b)–(c). Note that this new diagram does not include a representation of rod excitations. Previous studies have presented visualizations of rod excitation using traditional chromaticity diagrams [4]. In contrast, the present work focuses on visualizing the melanopsin signal. Rod and melanopsin signals exhibit a high degree of correlation, and no specific post-receptoral pathway is known to compare them directly. Therefore, representing either the rod or melanopsin signal allows for a three-dimensional depiction of the five-dimensional photoreceptor space, while still preserving its key structural features.

Because every axis is a photoreceptor signal divided by the same luminance term L + M, this is a linear space: an additive mixture of two lights plots on the straight line joining them, at a position set by their relative luminance. Any spectrum is a mixture of monochromatic lights, so it plots at the luminance-weighted combination of the corresponding points on the spectrum locus, which—being convex over most of its range—maps out the edge of the gamut, with all real-world spectra falling within it. These visualizations are therefore read like a gamut-coverage diagram: the real-world spectra map to a region, and a display is characterised by the subset of that region it can reproduce within tolerance. Good performance corresponds to reproducing the distribution of relevant real-world spectra as fully as possible. However, gamut coverage alone does not define the evaluation, because not all points in the physically attainable region are equally likely to occur. We therefore evaluate reproduction using the reference dataset described above, rather than assigning uniform weight to all possible points in the region.

## 3. Results: Example applications of novel metrics

To demonstrate the effectiveness of the proposed metrics, they are applied to three real three-primary displays, two hypothetical five-primary displays, and three research-grade five-primary displays.

Three real three-primary displays are evaluated: a CRT NEC MultiSync FP2141SB (NEC Display Solutions, Tokyo, Japan), a Dell U2715H LCD monitor (Dell, Austin, TX, USA), and a Display++ LCD monitor (Cambridge Research Systems, Rochester, UK). The spectral output of the display primaries was measured using a CRS SpectroCal MKII spectroradiometer (Cambridge Research Systems, Rochester, UK), as shown in Fig. 3(a)-(c). Two hypothetical five-primary displays are also evaluated: a hypothetical narrowband display, and a hypothetical broadband display (primaries depicted in Fig. 3(d)-(e)). We analyzed three custom-built research displays that use five or more primaries. These include a five-primary visual display unit (VDU) developed at the University of Manchester [34, 63], the multi-primary high dynamic range (RealVision MPHDR) display at the University of Oxford [62], and the five-primary Maxwellian-view display at Queensland University of Technology [65]. Figure 3(f)–(h) shows the primary spectra of the Manchester five-primary VDU, the RealVision MPHDR, and the Nugent–Zele display, respectively. The primaries in the hypothetical five-primary displays are either narrowband or broadband, with peak wavelengths at 450 nm, 500 nm, 550 nm, 600 nm, and 650 nm. The consequences of the primary placement are considered in the Discussion.

**Figure 3.**
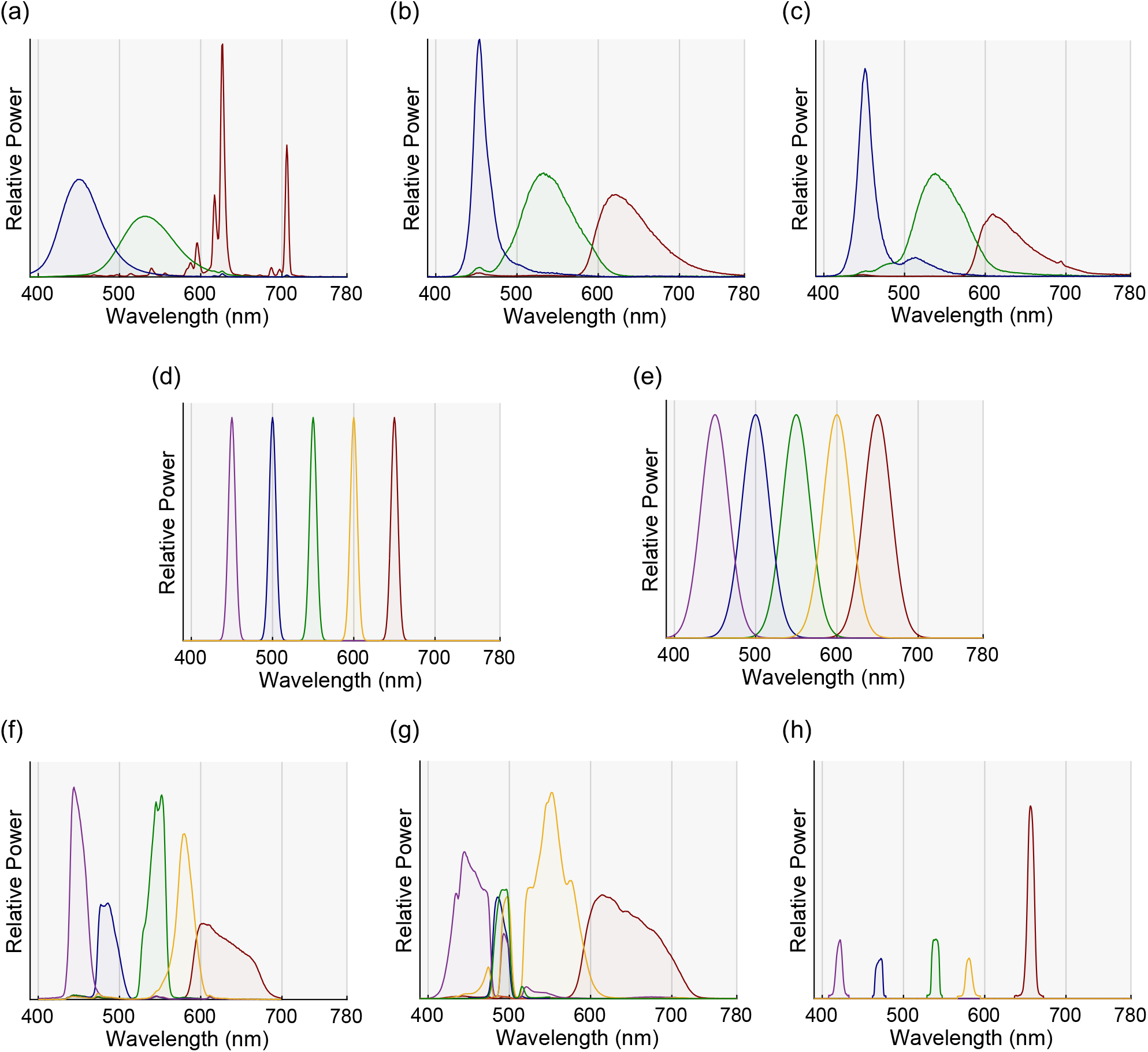
Display primaries used in the example applications of the proposed metrics. (a) CRT NEC Color Monitor. (b) Dell U2715H LCD Monitor. (c) CRS Display++ LCD Monitor. (d) Hypothetical narrowband five-primary display with 10 nm FWHM and peaks at 450, 500, 550, 600, and 650 nm. (e) Hypothetical broadband five-primary display with 40 nm FWHM and the same peak wavelengths. (f) Manchester five-primary VDU. (g) RealVision MPHDR display. (h) Nugent–Zele five-primary display.

### 3.1. Photoreceptor Signal Reproduction Metric (*PSRM*)

Most real-world signal chromaticities fall within the gamut of the selected displays, as shown in the CIE 1931 xy chromaticity diagram in Fig. 4(a). As shown in Fig. 4(b), each display can reproduce at least 90% of the chromaticities, specifically: CRT NEC Color Monitor, 91.0%; Dell U2715H LCD Monitor, 90.4%; CRS Display++ LCD Monitor, 90.2%; Manchester VDU, 98.8%; RealVision MPHDR, 97.3%; Nugent–Zele, 100.0%; and both the narrowband and broadband five-primary displays, 100%. Despite their excellent chromaticity reproduction, Fig. 4(c) shows that the three-primary displays have low *PSRM* values (CRT NEC Color Monitor: 2.5%; Dell U2715H LCD Monitor: 2.3%; CRS Display++ LCD Monitor: 0.7%). This is expected, as any match between the reproduced rod and melanopsin signals and those of real-world signals occurs by chance.

**Figure 4.**
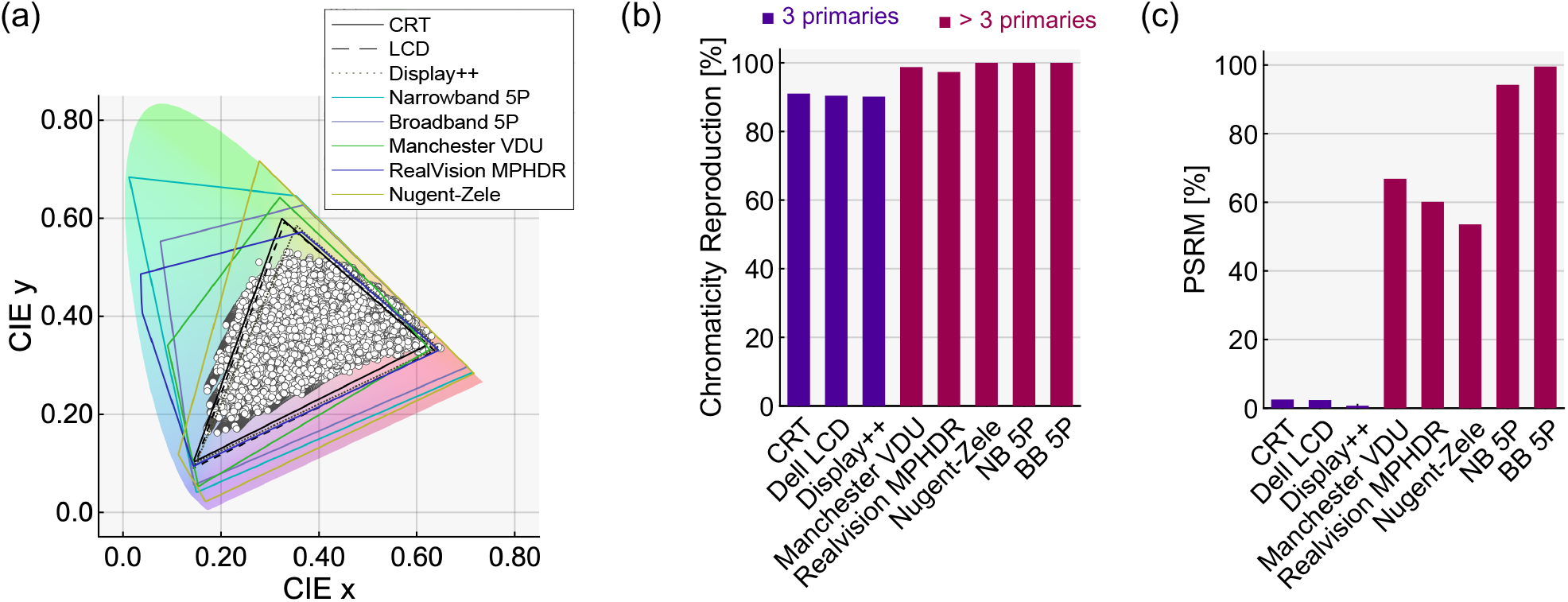
Chromaticity reproduction and *PSRM* of the real-world reference spectra across the example displays. (a) Chromaticities of 18,216 spectra plotted on the CIE 1931 xy chromaticity diagram, along with the color gamuts of the example displays. (b) Chromaticity reproduction rates of the example displays. (c) *PSRM* values of the example displays. While all displays show high chromaticity reproduction, *PSRM* is low for the three-primary displays and substantially higher for the five-primary displays. In panels (b) and (c), three-primary displays are shown in purple, and five-primary displays in pink.

Both hypothetical five-primary displays exhibit high *PSRM* values (Narrowband Five-Primary Display: 94.2%; Broadband Five-Primary Display: 99.5%). The variation in *PSRM* between the hypothetical five-primary displays indicates that, while five primaries are necessary to reproduce the α-opic quintuplet, not just any set of five will suffice. This is analogous to the fact that arbitrary selection of three primaries cannot guarantee a wide color gamut in three-primary displays. The *PSRM* values for the research-grade systems are lower than those of the hypothetical systems, with 66.8% for the Manchester five-primary VDU, 60.1% for the RealVision MPHDR, and 53.6% for the Nugent–Zele system. The relatively low *PSRM* of the real five-primary displays reflects their design focus on melanopsin contrast rather than reproducing real-world signals. This highlights that five or more primaries alone do not ensure a high *PSRM*; careful selection and optimization of primaries can improve reproduction accuracy. *PSRM* may serve as a useful metric for guiding primary selection in future multi-primary displays aimed at reproducing the real-world α-opic quintuplet.

### 3.2. Photoreceptor Signal Distortion Metric (*PSDM*)

Figure 5 illustrates rod and melanopsin signal distortions for three 3-primary displays: (a) NEC CRT, (b) Dell LCD, and (c) CRS Display++. Each panel plots the reproduced signal*A*_*ki*_’, against the real-world signal *A*_*ki*_, for each spectrum *i*, for *k* ∈ [*R, I*] (upper and lower plots, respectively), with cone signals matched. Deviations from the unity line indicate the extent of distortion between reproduced and real-world signals.

**Figure 5.**
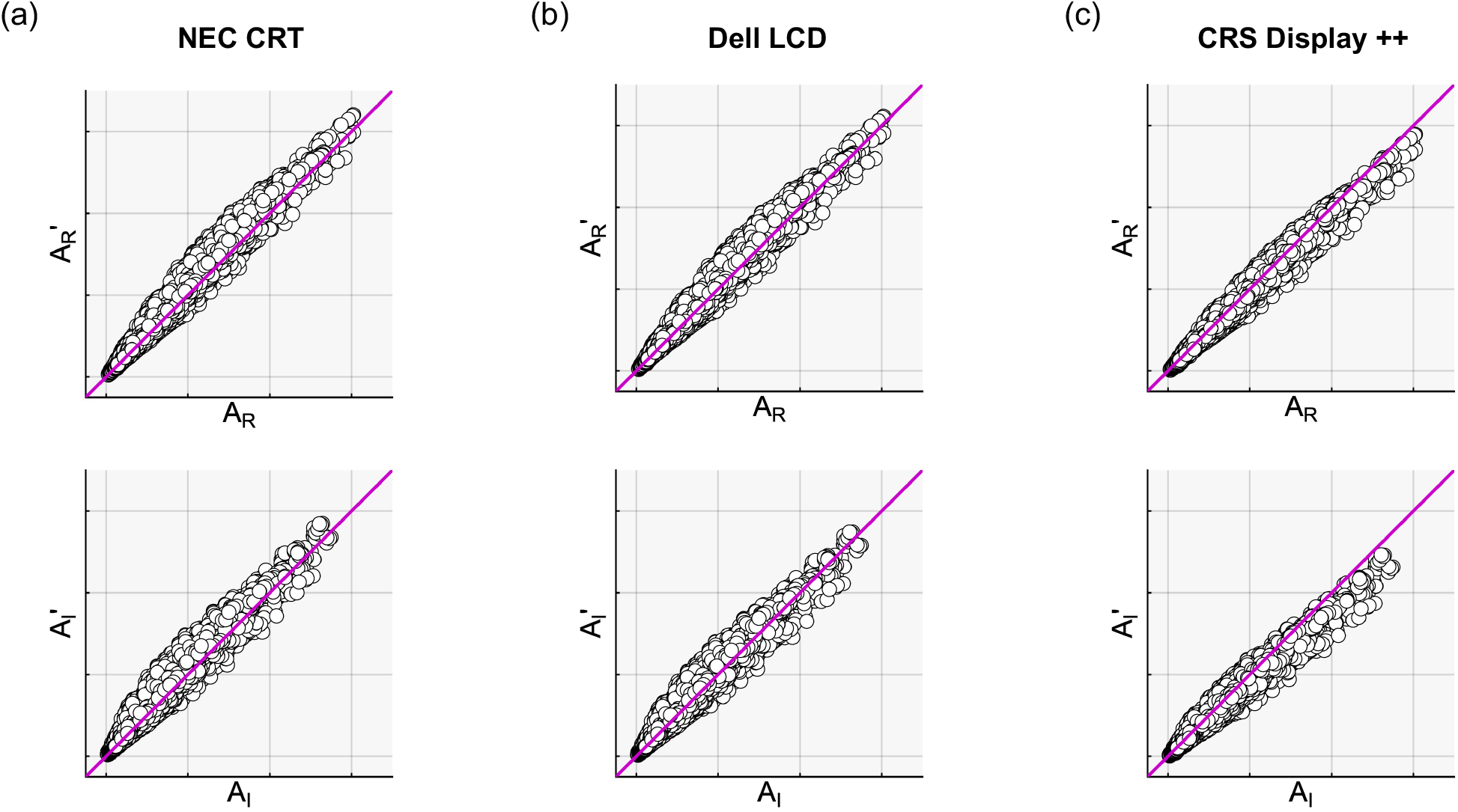
Reproduced rod (*k* = *R*) and melanopsin (*k* = *I*) signals *A*_*k*_*’* reproduced are plotted against their corresponding real-world signals *A*_*k*_ for each in-gamut spectrum across three types of three-primary displays: NEC CRT, Dell LCD, and CRS Display++ LCD.

The upper panels of Figure 6(a) show boxplots of rod *PSDM*_*ki*_ across individual spectra for three types of displays. To visualize chromaticity dependence, the CIE 1931 xy diagram is divided into 0.01 × 0.01 bins, and mean *PSDM*_*ki*_ values within each bin are averaged to produce *PSDM*_*kxy*_, shown in the lower panel of Fig. 6(a). For all three three-primary displays, rod *PSDM*_*ki*_ values were broadly distributed, with median signed *PSDM*_*Ri*_ of 2.6% for the CRT, −0.6% for the Dell LCD and −8.7% for the Display ++. The results for the melanopsin signal were similar (panel b), with median signed *PSDM*_*Ii*_ of 1.9% for the CRT, −2.0% for the Dell LCD and −13.5% for the Display ++. Overall, the Display++ showed the largest median signed distortions, strongly negative for both rod and melanopsin, indicating systematic under-reproduction of these signals relative to their real-world values; the CRT and Dell LCD showed smaller median distortions of mixed sign.

**Figure 6.**
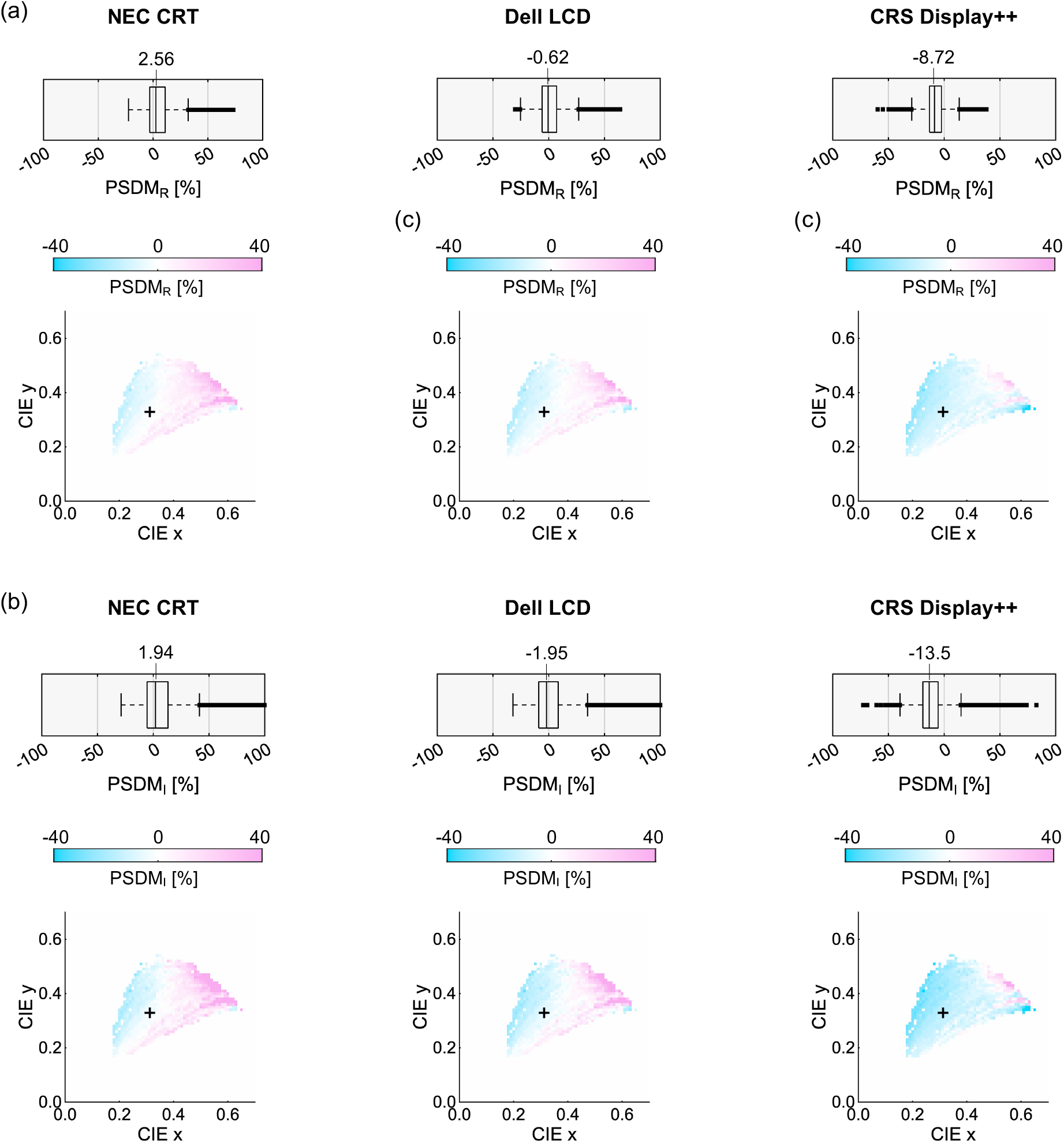
*PSDM* analysis applied to CRT, LCD and Display ++. (a) Rod signal analysis. Boxplots show the distribution of rod *PSDM*_*ki*_ across all individual spectra, and mean rod distortion across the CIE 1931 *xy* chromaticity diagram. For each 0.01 × 0.01 grid square, the average *PSDM*_*ki*_ of all spectra within that chromaticity bin is computed as *PSDM*_*kxy*_. The black plus shows the chromaticity of D65. (b) Melanopsin signal analysis. Same presentation as in panel (a).

### 3.3. Equal-luminance photoreceptor excitation visualizations

Figure 7(a) shows the 18,216 real-world spectra plotted in the proposed equal-luminance photoreceptor excitation diagram. The space is divided into a 100 × 100 grid. Spectra considered reproducible on each display (as defined by the *PSRM*) are shown in the proposed photoreceptor excitation space. This is illustrated in Fig. 7(b) for the three-primary displays and Fig. 8 for the five-primary displays.

**Figure 7.**
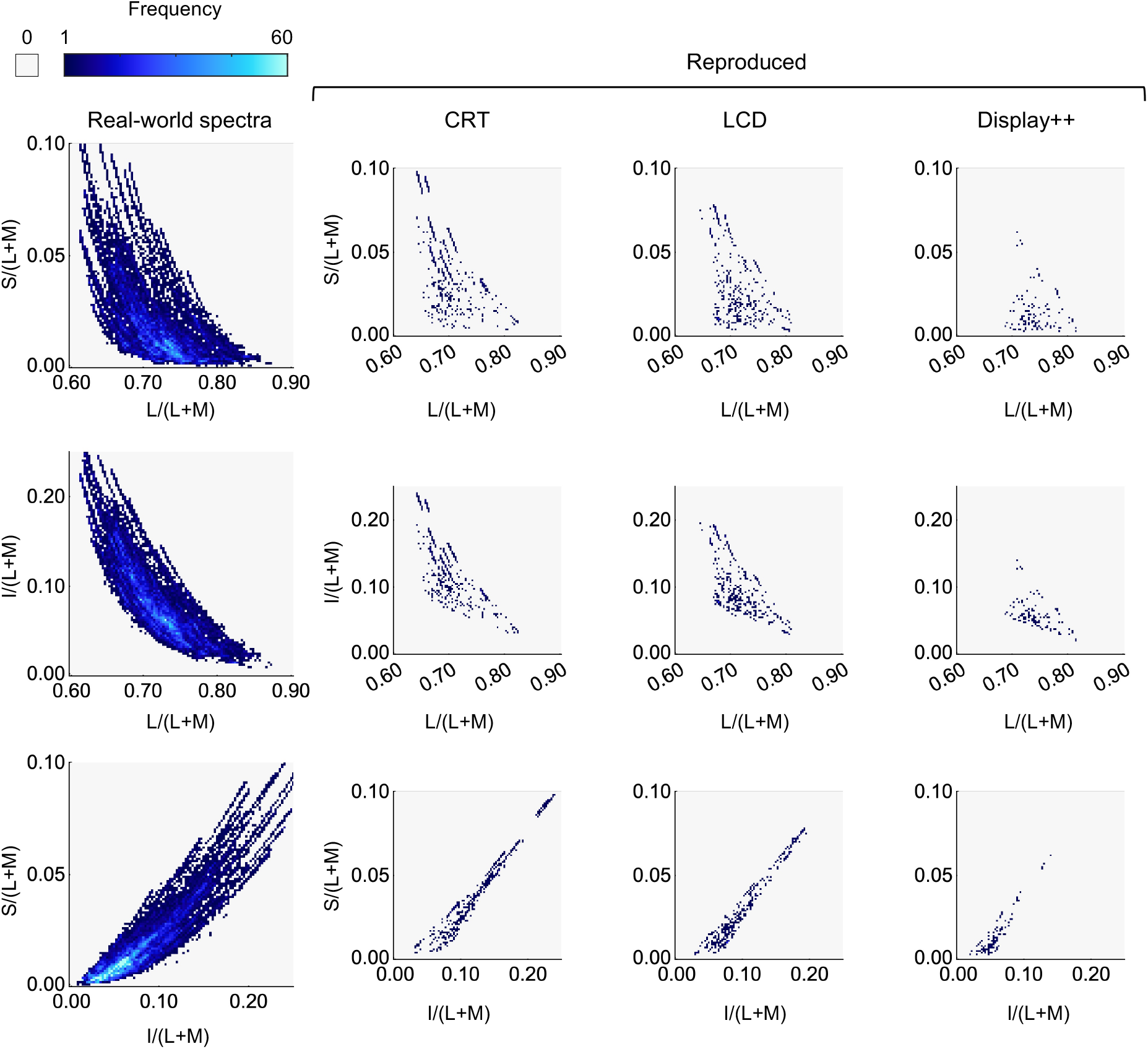
Visualization of α-opic signals from the real-world and the reproduced spectra in the proposed space. The leftmost column shows the real-world spectra plotted as a heat map, showing the number of spectra falling within each bin of a 100 × 100 grid. Other columns show spectra reproducible on example three-primary displays, where only spectra with distortion below the tolerance threshold (*ϵ* = 0.01) are included.

**Figure 8.**
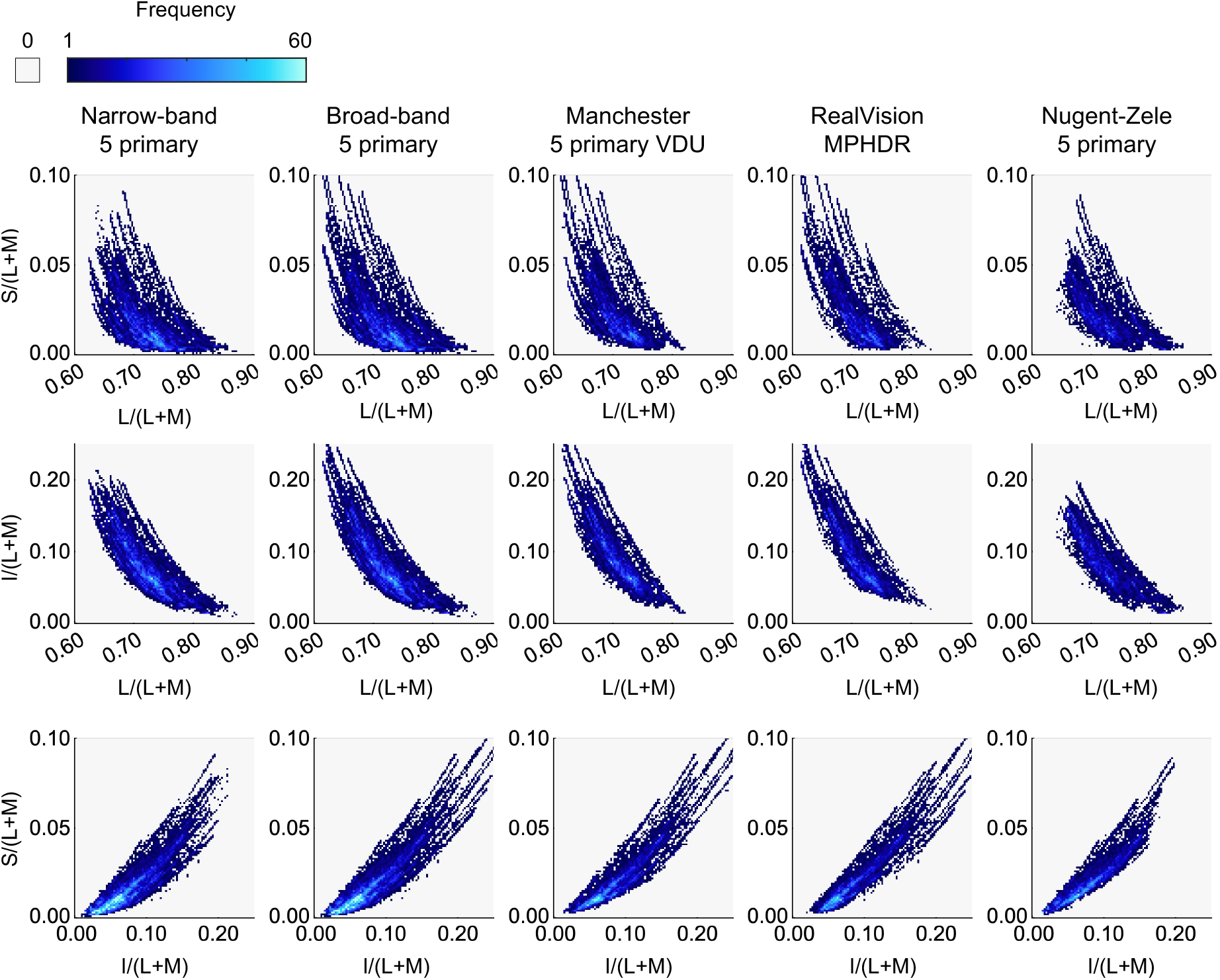
Reproduced spectra from the five-primary displays shown, using the same axes and color scale as in Fig. 7.

For the five-primary displays, the distribution of reproduced spectra closely resembles that of the real-world spectra in the proposed equal-luminance photoreceptor excitation space, though it covers a slightly smaller area due to imperfect reproduction (*PSRM* < 100%). In contrast, the three-primary displays produce a patchy and reduced distribution, highlighting their inability to fully reproduce all five real-world photoreceptor signals.

### 3.4. Robustness and additional analyses

The analyses above use the main reference dataset and the CIE S026 standard observer. Here, we assess the robustness of the metrics and the resulting conclusions to these choices, as well as to three additional factors: display bit depth, tolerance criteria, and the equal weighting assigned to the reference spectra.

#### 3.4.1. Reference dataset and real-world validation

The proposed metrics are not tied to any particular reference set; they can be applied to any collection of real-world spectra suits to the intended application. To test whether our conclusions depend on this choice, we re-evaluated every display on an independent set of directly measured real-world radiance spectra (the SpectroSense dataset [76]; 29,664 spectra), rather than on illuminant–reflectance products. The main findings were preserved (Table 1): the three-primary displays reproduced only a small fraction of the α-opic quintuplets (PSRM 2.2–6.5%), whereas the five-primary displays reproduced far more (72.1–86.2%), and the per-photoreceptor distortion pattern was unchanged, with the Display++ again showing the largest rod and melanopsin distortions. The absolute PSRM values differ from those on the main set, reflecting the different spectral composition of the two datasets, but the ordering of the displays and the central conclusion—that three primaries cannot reproduce the rod and melanopsin signals whereas well-placed five-primary displays can—are robust to the choice of reference set.

**Table 1.** Chromaticity reproduction and PSRM (%) on the main reference set (Main) and the directly measured SpectroSense dataset (SS).

| Display | Chrom. (Main) | PSRM (Main) | Chrom. (SS) | PSRM (SS) |
| --- | --- | --- | --- | --- |
| CRT NEC | 91.0 | 2.5 | 83.6 | 2.4 |
| Dell LCD | 90.4 | 2.3 | 83.6 | 6.5 |
| Display++ | 90.2 | 0.7 | 83.6 | 2.2 |
| Narrowband 5-primary | 100.0 | 94.2 | 90.6 | 82.2 |
| Broadband 5-primary | 100.0 | 99.5 | 89.7 | 86.2 |
| Manchester VDU | 98.8 | 66.8 | 84.1 | 85.2 |
| RealVision MPHDR | 97.3 | 60.1 | 83.8 | 72.1 |
| Nugent–Zeile | 100.0 | 53.6 | 95.7 | 85.9 |

#### 3.4.2. Individual variation in cone spectral sensitivities

Individual observers differ considerably in photoreceptor spectral sensitivity, even within the colour-normal population [77]; such differences underlie mean-observer metamerism, in which a display match for one observer is not a match for another [78]. One characteristic of the proposed framework is that individualised sensitivities can be substituted directly into the signal vectors and the MacLeod–Boynton axes [79, 80]. To quantify how much inter-observer variation affects the metrics, we generated 1,000 individual observers with the Asano individual-observer model, varying lens pigment density, macular pigment density, optical densities of L-, M-, and S-cone photopigments, and λ_max_ shifts of L-, M-, and S-cone photopigments [77]. Accordingly, this analysis assesses the combined influence of the sampled observer parameters and field size rather than variation at a fixed 10° field alone. Each observer’s cone fundamentals were propagated through the metrics. Inter-observer variation in lens optical density affects both rod and melanopsin sensitivities because light passes through the lens before reaching either photoreceptor. In contrast, macular pigment density affects rod sensitivity only, because melanopsin-containing retinal ganglion cells lie anterior to the macular pigment. We therefore reconstructed the rod and melanopsin sensitivity functions following the implementation in the Silent Substitution Toolbox [81], applying each observer’s lens density to both functions and their macular pigment density to the rod function alone. The rod and melanopsin photopigments were held at their standard values, as no population model of rod or melanopsin photopigment variation currently exists. Because the model samples observers around the population mean, the family-mean PSRM remained close to the standard-observer value for most displays (Table 2). Thus, the inter-observer spread is the more informative quantity. The exception was the RealVision MPHDR, for which the family-mean PSRM was lower than the standard-observer value. This spread (SD) was display-dependent: the broadband displays (the three-primary displays, the broadband five-primary display, and the Manchester VDU) remained stable, whereas the displays with narrowband primaries—the narrowband five-primary display, the RealVision MPHDR, and the Nugent–Zele system—were more variable. The per-photoreceptor distortions of the three-primary displays were likewise stable across observers (Table 3), indicating that the reported distortions are relatively robust to the modelled individual variations.

**Table 2.** Inter-observer variability of the PSRM (%): mean, standard deviation (SD), and 5–95% range across 1,000 individual observers. Rod and melanopsin sensitivities are reconstructed with the Silent Substitution Toolbox [81].

| Display | Mean | SD | 5–95% range |
| --- | --- | --- | --- |
| CRT NEC | 1.6 | 0.6 | 0.7–2.5 |
| Dell LCD | 1.7 | 0.7 | 0.8–3.1 |
| Display++ | 0.7 | 0.3 | 0.3–1.3 |
| Narrowband 5-primary | 94.9 | 6.1 | 82.7–99.7 |
| Broadband 5-primary | 98.2 | 1.7 | 95.7–99.6 |
| Manchester VDU | 65.9 | 5.0 | 56.8–73.1 |
| RealVision MPHDR | 27.0 | 15.7 | 9.0–52.0 |
| Nugent–Zeile | 51.3 | 10.3 | 30.8–65.4 |

**Table 3.** Inter-observer variability of the median signed PSDM_ki_ (%) for the three-primary displays: mean ± SD across 1,000 observers. Rod and melanopsin sensitivities carry each observer’s yoked lens variation (and, for the rod, macular pigment variation), reconstructed with the Silent Substitution Toolbox [81].

| Display | Rod (mean ± SD) | Melanopsin (mean ± SD) |
| --- | --- | --- |
| CRT NEC | 3.7 ± 0.9 | 2.5 ± 0.5 |
| Dell LCD | 0.3 ± 0.9 | -0.3 ± 1.9 |
| Display++ | -7.4 ± 1.0 | -12.7 ± 1.4 |

#### 3.4.3. Effect of display bit depth

The main analyses assume idealised displays with infinite bit depth, whereas real displays quantise each primary to a finite number of levels. We evaluated this for the Manchester VDU and the Nugent–Zele system, modelling each spectrum as reproduced at its fixed real-world intensity—the display does not rescale the overall intensity to use more of the code range—so the drive values are quantised at the display’s fixed full-scale resolution (the RealVision MPHDR was excluded because its six-primary, five-parameter control precludes a like-for-like comparison). Finite bit depth reduced the PSRM substantially (Fig. 9(a)): from 66.8% at infinite resolution to 47.0%, 15.7%, and 0.8% at 12-, 10-, and 8-bit for the Manchester VDU, and from 53.6% to 46.6%, 24.5%, and 3.4% for the Nugent–Zele system. Under this fixed-intensity normalisation, finite bit depth substantially reduced the PSRM, illustrating that quantisation can become important when target drive values occupy only a limited fraction of the available code range.

**Figure 9.**
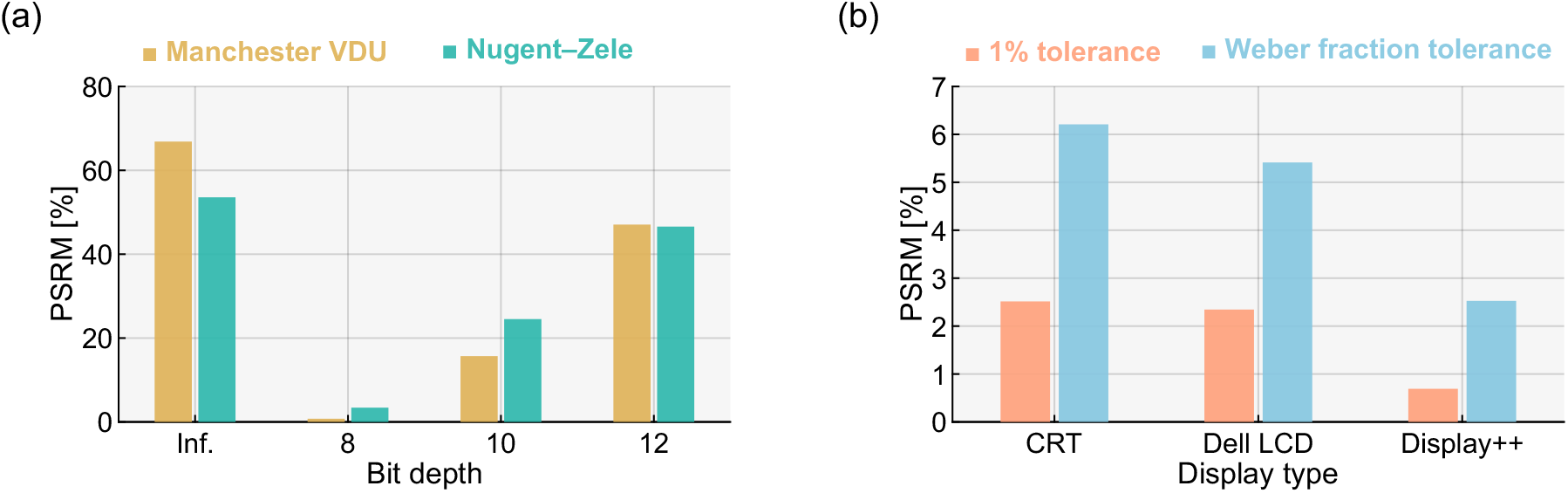
Influence of bit depth and tolerance value on *PSRM*. (a) Bar graphs show *PSRM* values for two five-primary display systems (Manchester VDU and Nugent–Zele system) across different bit depths. Infinity indicates the result from the original computation with no bit-depth limitation, while the other bars correspond to 8-, 10-, and 12-bit color depths commonly used in displays. (b) Effect of varying photoreceptor tolerance on *PSRM* across different display types.

#### 3.4.4. Effect of photoreceptor tolerance

The *PSRM* is inherently influenced by the tolerance thresholds applied. In the main analyses, a uniform 1% tolerance per photoreceptor was used. However, cone photoreceptor sensitivity—i.e., the increment thresholds for color discrimination—varies by type. We also examined receptor-specific tolerances based on reported Weber fractions: 2% for L and M cones, 9% for S cones, 14% for rods, and 1% for melanopsin [82–84]. Applying these values results in no change in *PSRM* for any of the five-primary displays, as exact reproduction across all five signals is already met. For three-primary displays, the relaxed rod tolerance leads to slight increases in *PSRM* (Figure 9(b)).

#### 3.4.5. Effect of luminance weighting

The metrics weight all reference spectra equally. However, in some cases, it may be preferable to assign greater weight to brighter samples, as they are more likely to attract attention. We therefore recomputed the metrics with each spectrum weighted by its photopic luminance (normalised to a maximum of one). Across all eight displays the PSRM changed by at most 4.8 percentage points (the Manchester VDU, 66.8→71.6%) and the overall PSDM by at most 0.6, indicating that the principal conclusions were insensitive to this particular relative photopic-luminance weighting scheme. Because the illuminants were normalised to unit area, this analysis does not evaluate weighting by absolute real-world scene luminance.

## 4. Discussion

Until now, display reproduction metrics have largely focused on cone signal fidelity (with some exceptions, e.g., Shapiro et al., 1996). While colour gamuts are a long-standing measure, they are inherently limited to describing chromaticity by design. Growing evidence that melanopsin contributes to non-visual (and possibly visual) responses, along with the possibility that rods influence vision across a broader range of light levels than previously assumed, has raised interest in extending display considerations to include rod and melanopsin reproduction. To support optimization, there is a need for metrics that quantify a display’s ability to reproduce—or, where direct control is lacking, the extent to which it distorts—the α-opic quintuplet. This paper proposed two such metrics.

### 4.1. The applicability of the proposed metrics

The *PSRM* quantifies how many α-opic quintuplets from a reference dataset can be reproduced without distortion, such that the original signal ratios are preserved. It is expressed as a single value for any display, reflecting its reproduction capability. The *PSRM* is analogous to the gamut metrics that are widely used for three-primary displays. The *PSDM* measures the distortion introduced when reproducing α-opic quintuplets on displays with fewer than five primaries. *PSDM* can be reported in several ways: as an overall average across all photoreceptors and spectra, or in a more detailed form per photoreceptor k and spectrum *i* (*PSDM*_*ki*_). A useful approach may be to compute the overall *PSDM* as a single value and to visualize the *PSDM*_ki_ as shown in Figures 5 and 6. Equal-luminance photoreceptor excitation diagrams Figures 7 and 8 are proposed as a means to visualize where display reproduction succeeds and where distortions may occur.

While there is no direct mathematical relationship between these metrics and existing color rendering metrics, useful analogies exist. Since the *PSRM* is based on reproduction of a reference set of spectra, it is conceptually similar to the Colour Rendering Index (CRI), which evaluates how well a light source reproduces reflectances under a reference illuminant. However, unlike CRI, which averages graded scores (0–100) across samples, *PSRM* is a binary count, effectively treating each sample as either fully reproduced or not, then expressing the result as a percentage. The *PSDM*, in turn, is analogous to the *ΔE* color difference metric, but quantifies reproduction error in terms of photoreceptor signals rather than distances in CIELAB color space.

The metrics and visualization proposed here aim to provide a useful framework for characterizing an image reproduction system’s ability to accurately reproduce the α-opic quintuplet from real-world scenes. As understanding grows around the roles of rod and melanopsin signals in both visual and non-visual responses to light, interest in display metrics that account for these photoreceptors is increasing, particularly in applications aiming to replicate naturalistic visual experiences or support emerging aspects of visual function, such as virtual and augmented reality, the metaverse, 3D displays, and advanced “night-shift” modes. The proposed metrics are intended to support this evolving perspective by offering tools to assess and visualize photoreceptor signal reproduction beyond the cones.

### 4.2. Limitations of proposed metrics

The metrics introduced here quantify how well retinal-level α-opic signals are reproduced, without making assumptions about the downstream perceptual, circadian, or cognitive interpretation of those signals. Thus, they should be understood as measures of low-level physiological fidelity rather than direct predictors of higher-level outcomes. While one could envision mechanistic models that integrate photoreceptor signals to predict visual and non-visual effects of light exposure [85], further research is needed to understand how these signals combine to drive specific responses. For example, the relationship between varying levels of melanopsin signal distortion and melatonin suppression at different circadian phases remains to be quantified. More generally, determining the perceptual and non-visual consequences of accurately reproduced rod and melanopsin signals remains an open question.

Given the increasing proportion of time that users spend viewing displays [86] and given the known inputs from retinal photoreceptors to other aspects of our physiology, such as the circadian system, quantification of the mismatch between real-world exposure and screen exposure is becoming increasingly important. Whilst we recognise that content (e.g. characteristics of the real-environment, and whether screen content is video-recording, graphics or text) is an important part of this picture (not investigated here), the proposed metrics raise awareness of the issue for direct reproduction of real-world spectra.

A further limitation is the disregard for absolute intensity. This could be addressed by requiring the *PSRM* to match both the spectral and intensity profiles of real-world α-opic quintuplets. Alternatively, metrics could be reported with the display’s intensity range or limited to applicable intensity levels. A major challenge is defining the typical intensity range of real-world spectra, which relates to the broader need to characterize the human “spectral diet” [72]. Furthermore, capturing intensity is complex due to its dependence on lighting [87] and viewing geometry [88].

The real-world reference dataset used here was selected because it underpins the development of the IES TM-30 colour fidelity method [73] and its 99 colour-evaluation samples are used in the IES TM-30 standard. However, it excludes structural and interference colours, which though less common, are perceptually salient, and does not weight samples by the likelihood of encountering them in real-world settings [52, 72].

### 4.3. Applying the metrics and visualization to multi-primary displays

Having sufficient primaries to control five photoreceptors does not ensure a high *PSRM*, just as having any three primaries does not guarantee a wide chromaticity gamut. This is shown by the substantially lower PSRM of the research-grade five-primary systems (53–67%) and by the gap between the two hypothetical displays, where the narrowband display (94.2%) falls short of the broadband display (99.5%) despite sharing the same primary wavelengths. Achieving high *PSRM* requires careful optimization of the primaries, but this is not always the goal. Many five-primary systems, including the two custom-built ones described here, were designed for specific research purposes and use primaries suited to those aims, which can result in lower *PSRM*. The same applies to three-primary displays, which are not designed to reproduce rod and melanopsin signals. As a result, they show high *PSRM* for cone-based chromaticity reproduction but low *PSRM* across all five photoreceptors. Moreover, a limited number of bits per channel can also introduce a significant reduction in PSRM (see section 3.4.3).

Systems with five or more primaries can control all five photoreceptor signals and, when driven to reproduce the α-opic quintuplet, have a *PSDM* of zero for in-gamut signals. *PSDM* is therefore mainly relevant for displays with fewer than five primaries or for five-primary systems not developed to control all five signals. An exception is evaluating tone mapping, where *PSDM* can quantify distortions from mapping out-of-gamut signals into gamut for five-primary systems. Thus, the visualizations in Figures 5 and 6 apply only when *PSDM* is non-zero and are not shown for the five-primary displays. In contrast, the extended MacLeod-Boynton diagram (Figure 7) applies to all displays and can be plotted like a chromaticity gamut to visualize the full α-opic space. A similar visualization can also be achieved by extending other chromaticity diagrams, such as the CIE xy diagram [89].

The proposed metrics may also aid in designing four-primary displays [90], which can match four of five photoreceptor signals. Given the strong correlation between rod and melanopsin responses, a practical trade-off may be achievable, e.g. targeting cone and melanopsin control could yield low *PSDM* for rod signals and high *PSRM*. These metrics could therefore guide the selection of primaries and control strategies to optimize *PSRM* while minimizing *PSDM*.

## 5. Conclusion

This study introduces two physiologically relevant display metrics—the photoreceptor signal reproduction metric (*PSRM*) and the photoreceptor signal distortion metric (*PSDM*)—to characterize the full photoreceptor response to light from displays. It also proposes a new visualization tool: an equal-luminance photoreceptor excitation diagram, extending the MacLeod-Boynton chromaticity diagram. These methods enable quantification of signals beyond the cones, supporting evaluation and design of multi-primary displays for accurate reproduction of photoreceptor responses. The metrics also help characterize distortions introduced by displays with fewer than five primaries or by five-primary systems not controlling the full α-opic quintuplet. While not intended as definitive, these tools offer a useful foundation for incorporating currently overlooked photoreceptor signals into future display evaluation frameworks.

## Acknowledgements

The authors are grateful to Thomas Nugent and Andrew Zele for providing the spectra of the primaries of their display [65].

## Data availability

The data underlying the results presented in this paper and the code to apply the novel metrics and visualization (written in MATLAB R2025b (Mathworks, Natick, MA, USA)) are publicly available at https://github.com/takuma929/metrics_photoreceptor_signal_reproduction and Zenodo repository (released at the time of publication).

## References

CIE, “CIE System for Metrology of Optical Radiation for ipRGC-Influenced Responses to Light. CIE S 026/E:2018,” (Vienna: Central Bureau of the Commission Internationale de l’Éclairage) (2018).

D. I. A. MacLeod and R. M. Boynton, “Chromaticity diagram showing cone excitation by stimuli of equal luminance,” J. Opt. Soc. Am. 69, no. 8 (1979): 1183–1186.

M. Aguilar and W. Stiles, “Saturation of the rod mechanism of the retina at high levels of stimulation,” Opt. Acta: Int. J. Opt. 1, no. 1 (1954): 59–65.

A. G. Shapiro, J. Pokorny, and V. C. Smith, “Cone-rod receptor spaces with illustrations that use CRT phosphor and light-emitting-diode spectra,” J. Opt. Soc. Am. A 13, no. 12 (1996): 2319–2328.

L. T. Sharpe and A. Stockman, “Rod pathways: the importance of seeing nothing,” Trends Neurosci. 22, no. 11 (1999): 497–504.

G. S. Lall, V. L. Revell, H. Momĳi, J. A. Enezi, C. M. Altimus, A. D. Güler, C. Aguilar, M. A. Cameron, S. Allender, M. W. Hankins, and R. J. Lucas, “Distinct contributions of rod, cone, and melanopsin photoreceptors to encoding irradiance,” Neuron 66 (2010): 417–428.

R. Frederiksen, A. Morshedian, S. A. Tripathy, T. Xu, G. H. Travis, G. L. Fain, and A. P. Sampath, “Rod photoreceptors avoid saturation in bright light by the movement of the g protein transducin,” J. Neurosci. 41 (2021): 3320–3330.

I. C. Méndez, A. Martín, B. O’Donell, D. Cao, and P. A. Barrionuevo, “Temporal integration of rod signals in luminance and chromatic pathways,” JOSA A 39 (2022): 1782–1793.

S. Uprety, P. Adhikari, B. Feigl, and A. J. Zele, “Melanopsin photoreception differentially modulates rod-mediated and cone-mediated human temporal vision,” iScience 25 (2022).

A. J. Zele and D. Cao, “Vision under mesopic and scotopic illumination,” Front. Psychol. 5 (2015): 1584.

A. Stockman and L. T. Sharpe, “Into the twilight zone: the complexities of mesopic vision and luminous efficiency,” Ophthalmic Physiol. Opt. 26, no. 3 (2006): 225–239.

F. Zaidi, J. Hull, S. Peirson, K. Wulff, D. Aeschbach, J. Gooley, G. Brainard, K. Gregory-Evans, J. Rizzo, C. Czeisler, R. Foster, M. Moseley, and S. Lockley, “Short-wavelength light sensitivity of circadian, pupillary, and visual awareness in humans lacking an outer retina,” Curr. Biol. 17, no. 24 (2007): 2122–2128.

A. Zele, B. Feigl, P. Adhikari, M. Maynard, and D. Cao, “Melanopsin photoreception contributes to human visual detection, temporal and colour processing,” Sci. Reports 8 (2018): 3842.

M. Spitschan, A. S. Bock, J. Ryan, G. Frazzetta, D. H. Brainard, and G. K. Aguirre, “The human visual cortex response to melanopsin-directed stimulation is accompanied by a distinct perceptual experience,” Proc. Natl. Acad. Sci. 114, no. 46 (2017): 12291–12296.

D. Cao, A. Chang, and S. Gai, “Evidence for an impact of melanopsin activation on unique white perception,” J. Opt. Soc. Am. A 35, no. 4 (2018): B287–B291.

T. M. Brown, S. I. Tsujimura, A. E. Allen, J. Wynne, R. Bedford, G. Vickery, A. Vugler, and R. J. Lucas, “Melanopsin-based brightness discrimination in mice and humans,” Curr. Biol. 22, no. 12 (2012): 1134–1141.

A. J. Zele, P. Adhikari, B. Feigl, and D. Cao, “Cone and melanopsin contributions to human brightness estimation,” J. Opt. Soc. Am. A 35 (2018): B19–B25.

S.-E. Chien, S.-L. Yeh, W. Yamashita, and S.-i. Tsujimura, “Enhanced human contrast sensitivity with increased stimulation of melanopsin in intrinsically photosensitive retinal ganglion cells,” Vis. Res. 209 (2023): 108271.

P. A. Barrionuevo, M. L. S. Salinas, and J. M. Fanchini, “Are ipRGCs involved in human color vision? Hints from physiology, psychophysics, and natural image statistics,” Vis. Res. 217 (2024): 108378.

P. A. Barrionuevo, C. P. Filgueira, and D. Cao, “Is melanopsin activation affecting large field color-matching functions?” JOSA A 39 (2022): 1104–1110.

T. Woelders, A. E. Allen, and R. J. Lucas, “Melanopsin enhances image persistence,” Curr. Biol. 33 (2023): 5048–5056.

T. DeLawyer and K. Shinomori, “Melanopsin-driven surround induction on the red/green balance of yellow,” JOSA A 40 (2023): A40–A47.

T. M. Brown, “Melanopic illuminance defines the magnitude of human circadian light responses under a wide range of conditions,” J. Pineal Res. 69, no. 1 (2020): e12655.

J. F. Duffy and C. A. Czeisler, “Effect of light on human circadian physiology,” Sleep Medicine Clin. 4, no. 2 (2009): 165–177.

C. Blume, C. Garbazza, and M. Spitschan, “Effects of light on human circadian rhythms, sleep and mood,” Somnologie 23, no. 3 (2019): 147–156.

R. J. Lucas, S. N. Peirson, D. M. Berson, T. M. Brown, H. M. Cooper, C. A. Czeisler, M. G. Figueiro, P. D. Gamlin, S. W. Lockley, J.B. O’Hagan et al., “Measuring and using light in the melanopsin age,” Trends Neurosci. 37 (2014): 1–9.

G. C. Brainard, J. P. Hanifin, J. M. Greeson, B. Byrne, G. Glickman, E. Gerner, and M. D. Rollag, “Action spectrum for melatonin regulation in humans: Evidence for a novel circadian photoreceptor,” J. Neurosci. 21, no. 16 (2001): 6405–6412.

D. Dacey, H. Liao, B. Peterson, R. Farel, V. Smith, J. Pokorny, K. Yau, and P. Gamlin, “Melanopsin-expressing ganglion cells in primate retina signal colour and irradiance and project to the LGN,” Nature 433, no. 7027 (2005): 749–754.

P. Gamlin, D. McDougal, J. Pokorny, V. Smith, K. Yau, and D. Dacey, “Human and macaque pupil responses driven by melanopsin-containing retinal ganglion cells,” Vis. Res. 47, no. 7 (2007): 946–954.

P. A. Barrionuevo and D. Cao, “Luminance and chromatic signals interact differently with melanopsin activation to control the pupil light response,” J. Vis. 16, no. 11 (2016): 29.

D. Cao, N. Nicandro, and P. A. Barrionuevo, “A five-primary photostimulator suitable for studying intrinsically photosensitive retinal ganglion cell functions in humans,” J. Vis. 15, no. 1 (2015): 15.1.27.

S. Tsujimura, K. Ukai, D. Ohama, A. Nuruki, and K. Yunokuchi, “Contribution of human melanopsin retinal ganglion cells to steady-state pupil responses,” Proc. Royal Soc. B Biol. Sci. 277 (2010): 2485–2492.

S. Tsujimura and Y. Tokuda, “Delayed response of human melanopsin retinal ganglion cells on the pupillary light reflex,” Ophthalmic & Physiol. Opt. 31, no. 5 (2011): 469–479.

M. Spitschan, M. Gardasevic, F. Martial, R. Lucas, and A. Allen, “Pupil responses to hidden photoreceptor-specific modulations in movies,” PLOS One 14, no. 5 (2019): e0216307.

P. A. Barrionuevo, L. A. Issolio, and C. Tripolone, “Photoreceptor contributions to the human pupil light reflex,” J. Photochem. Photobiol. 15 (2023): 100178.

M. Spitschan, S. Jain, D. H. Brainard, and G. K. Aguirre, “Opponent melanopsin and S-cone signals in the human pupillary light response,” Proc. Natl. Acad. Sci. 111, no. 43 (2014): 15568–15572.

M. Spitschan, “Photoreceptor inputs to pupil control,” J. Vis. 19, no. 9 (2019): 5, 1–5.

R. Lok, K. C. Smolders, D. G. Beersma, and Y. A. de Kort, “Light, alertness, and alerting effects of white light: A literature overview,” J. Biol. Rhythm. 33, no. 6 (2018): 589–601.

J. L. Souman, T. Borra, I. de Goĳer, L. J. Schlangen, B. N. Vlaskamp, and M. P. Lucassen, “Spectral tuning of white light allows for strong reduction in melatonin suppression without changing illumination level or color temperature,” J. Biol. Rhythm. 33, no. 4 (2018): 420–431.

C. Cajochen, “Alerting effects of light,” Sleep Medicine Rev. 11, no. 6 (2007): 453–464.

N. Milosavljevic, “How does light regulate mood and behavioral state?” Clocks & Sleep 1 (2019): 319–331.

A. Stockman, L. Sharpe, and C. Fach, “The spectral sensitivity of the human short-wavelength cones,” Vis. Res. 39, no. 17 (1999): 2901–2927.

A. Stockman and L. Sharpe, “The spectral sensitivities of the middle- and long-wavelength-sensitive cones derived from measurements in observers of known genotype,” Vis. Res. 40, no. 13 (2000): 1711–1737.

CIE, “Fundamental chromaticity diagram with physiological axes. Part 1. Technical Report 170-1. CIE 170-1:2006,” (Vienna: Central Bureau of the Commission Internationale de l’Éclairage) (2006).

G. Wyszecki and W. Stiles, Color Science: Concepts and Methods, Quantitative Data and Formulae, 2nd ed. (Wiley, 1982).

L.-H. Lee, T. Braud, P. Zhou, L. Wang, D. Xu, Z. Lin, A. Kumar, C. Bermejo, and P. Hui, “All one needs to know about metaverse: A complete survey on technological singularity, virtual ecosystem, and research agenda,” arXiv preprint 2110.05352 (2021).

“Mark in the Metaverse,” https://www.theverge.com/22588022/mark-zuckerberg-facebook-ceo-metaverse-interview. Accessed: 2023-05-26.

J. Arendt, “Melatonin and human rhythms,” Chronobiol. Int. 23 (2006): 21–37.

A. W. Wood, S. P. Loughran, and C. Stough, “Does evening exposure to mobile phone radiation affect subsequent melatonin production?” Int. J. Radiat. Biol. 82 (2006): 69–76.

M. G. Figueiro, B. Wood, B. Plitnick, and M. S. Rea, “The impact of light from computer monitors on melatonin levels in college students,” Neuroendocrinol. Lett. 32 (2011): 158–163.

C. Cajochen, S. Frey, D. Anders, J. Späti, M. Bues, A. Pross, R. Mager, A. Wirz-Justice, and O. Stefani, “Evening exposure to a light-emitting diodes (LED)-backlit computer screen affects circadian physiology and cognitive performance,” J. Appl. Physiol. 110, no. 5 (2011): 1432–1438.

B. Wood, M. S. Rea, B. Plitnick, and M. G. Figueiro, “Light level and duration of exposure determine the impact of self-luminous tablets on melatonin suppression,” Appl. Ergonomics 44 (2013): 237–240.

L. A. Ostrin, K. S. Abbott, and H. M. Queener, “Attenuation of short wavelengths alters sleep and the ipRGC pupil response,” Ophthalmic Physiol. Opt. 37 (2017): 440–450.

S. Cain, E. McGlashan, P. Vidafar, J. Mustafovska, S. Curran, X. Wang, A. Mohamed, V. Kalavally, and A. Phillips, “Evening home lighting adversely impacts the circadian system and sleep,” Sci. Reports 10 (2020): 19110.

W. Nagel, “Appendix: adaptation, twilight vision and the duplicity theory,” in Helmholtz’s Treatise on Physiological Optics, transl. J. P. C. Southall (1924): 313–343.

J. Pokorny, M. Lutze, D. Cao, and A. J. Zele, “The color of night: Surface color perception under dim illuminations,” Vis. Neurosci. 23 (2006): 525–530.

J. Pokorny, M. Lutze, D. Cao, and A. J. Zele, “The color of night: Surface color categorization by color defective observers under dim illuminations,” Vis. Neurosci. 25 (2008): 475–480.

S. L. Elliott and D. Cao, “Scotopic hue percepts in natural scenes,” J. Vis. 13 (2013): 15–15.

R. Nagare, M. S. Rea, B. Plitnick, and M. G. Figueiro, “Effect of white light devoid of ‘cyan’ spectrum radiation on nighttime melatonin suppression over a 1-h exposure duration,” J. Biol. Rhythm. 34, no. 2 (2019): 195–204.

S. L. Chellappa, C. J. Morris, and F. A. Scheer, “Circadian misalignment increases mood vulnerability in simulated shift work,” Sci. Reports 10 (2020): 1–10.

S. Ravikumar, K. Akeley, and M. S. Banks, “Creating effective focus cues in multi-plane 3D displays,” Opt. Express 19, no. 21 (2011): 20940–20952.

A. C. Hexley, A. Özgür Yöntem, M. Spitschan, H. E. Smithson, and R. Mantiuk, “Demonstrating a multi-primary high dynamic range display system for vision experiments,” J. Opt. Soc. Am. A 37, no. 4 (2020): A271–A284.

A. E. Allen, F. P. Martial, and R. J. Lucas, “Form vision from melanopsin in humans,” Nat. Commun. 10 (2019): 2274.

P. A. Barrionuevo, O. U. Preciado, M. L. Sandoval Salinas, and L. A. Issolio, “Chapter 2 - Optical stimulation systems for studying human vision,” in Circadian and Visual Neuroscience, vol. 273 of Progress in Brain Research, N. Santhi and M. Spitschan, eds. (Elsevier, 2022): 13–36.

T. W. Nugent and A. J. Zele, “A five-primary Maxwellian-view display for independent control of melanopsin, rhodopsin, and three-cone opsins on a fine spatial scale,” J. Vis. 22 (2022): 20–20.

T. W. Nugent, D. D. Carter, S. Uprety, P. Adhikari, B. Feigl, and A. J. Zele, “Protocol for isolation of melanopsin and rhodopsin in the human eye using silent substitution,” STAR Protocols 4 (2023): 102126.

CIE, “Commission Internationale de l’Éclairage Proceedings, 1931,” (Cambridge: Cambridge University Press, 1932).

CIE, “Fundamental chromaticity diagram with physiological axes. Part 2. Technical Report 170-2. CIE 170-2:2015,” (Vienna: Central Bureau of the Commission Internationale de l’Éclairage) (2015).

International Electrotechnical Commission, “IEC 61966-2-1:1999,” (1999).

Adobe Systems Incorporated, “Adobe RGB (1998) Color Image Encoding,” (2005).

International Telecommunications Union, “ITU-R BT.709-6: Parameter values for the HDTV standards for production and international programme exchange,” (2015).

F. S. Webler, M. Spitschan, R. G. Foster, M. Andersen, and S. N. Peirson, “What is the ‘spectral diet’ of humans?” Curr. Opin. Behav. Sci. 30 (2019): 80–86.

A. David, P. T. Fini, K. W. Houser, Y. Ohno, M. P. Royer, K. A. G. Smet, M. Wei, and L. Whitehead, “Development of the IES method for evaluating the color rendition of light sources,” Opt. Express 23, no. 12 (2015): 15888–15906.

K. Houser, M. Wei, A. David, M. Kremers, and X. Shen, “Review of measures for light-source color rendition and considerations for a two-measure system for characterizing color rendition,” Opt. Express 21, no. 8 (2013): 10393–10411.

Illuminating Engineering Society, ANSI/IES TM-30-18: IES Method for Evaluating Light Source Color Rendition, Illuminating Engineering Society, New York, 2018.

R. Lazar, J. Degen, A. S. Fiechter, A. Monticelli, and M. Spitschan, “Regulation of pupil size in natural vision across the human lifespan,” R. Soc. Open Sci. 11, 6 (2024).

Y. Asano, M. Fairchild, and L. Blondé, “Individual Colorimetric Observer Model,” PLOS One 11, no. 2 (2016): e0145671.

M. D. Fairchild and D. R. Wyble, “Mean observer metamerism and the selection of display primaries,” Proc. IS&T 15th Color and Imaging Conf., 151–156 (2007).

A. Stockman and A. T. Rider, “Formulae for generating standard and individual human cone spectral sensitivities,” Color Res. Appl. 48, 818–840 (2023).

A. C. Schneider, T. Morimoto, M. McClements, R. E. MacLaren, and H. E. Smithson, “Investigating ethnicity-related variability in the human L-cone spectral sensitivity function,” Vision Research, 243, 108769 (2026). 10.1016/j.visres.2026.108769.

M. Spitschan, G. K. Aguirre, and D. H. Brainard, “Selective stimulation of penumbral cones reveals perception in the shadow of retinal blood vessels,” PLOS One 10, no. 4 (2015): e0124328.

T. Cornsweet, Visual Perception (Academic Press, 2012).

H. Davson, Physiology of the Eye (Bloomsbury Publishing, 1990).

W. Stiles, “Color vision: The approach through increment-threshold sensitivity,” (1959).

A. C. Skeldon, T. R. Garcia, S. F. Cleator, C. della Monica, K. K. Ravindran, V. L. Revell, and D.-J. Dĳk, “Method to determine whether sleep phenotypes are driven by endogenous circadian rhythms or environmental light by combining longitudinal data and personalised mathematical models,” bioRxiv (2023).

D. L. Harvey, K. Milton, A. P. Jones, and A. J. Atkin, “International trends in screen-based behaviours from 2012 to 2019,” Prev. Med. 154 (2022): 106909.

A. Ryer, U. Light, and V. Light, Light Measurement Handbook (Citeseer, 1997).

T. Morimoto, S. Kishigami, J. M. Linhares, S. M. Nascimento, and H. E. Smithson, “Hyperspectral environmental illumination maps: Characterizing directional spectral variation in natural environments,” Opt. Express 27 (2019): 32277–32293.

G. Evéquoz, F. Truffer, and M. Geiser, “Maximum possible contrast level for silent substitution: a theoretical model applied to melanopsin stimulation,” J. Opt. Soc. Am. A 38, 1312–1319 (2021).

J. Pokorny, H. Smithson, and J. Quinlan, “Photostimulator allowing independent control of rods and the three cone types,” Vis. Neurosci. 21, no. 3 (2004): 263–267.

